# Engineered Miniature ParTasR Enables Efficient PAMless Genome Editing

**DOI:** 10.64898/2026.11.12.766130

**Authors:** Qianqian Zhu, Ziyi Zhang, Qiuyu Cao, Xiaojie Ma, Yuqiang Zeng, Mengyu Shang, Xiaohui Zhang

## Abstract

TIGR-Tas system holds great promise for genome-engineering applications owing to its compact size and PAMless targeting capability. However, its low editing efficiency in mammalian cells severely limits its broader utility. Here, we engineered a highly active variant of ParTasR, termed eTasR, through structure-guided rational design of ParTasR combined with co-expression of the 3′→5′ exonuclease TREX2. eTasR increased editing efficiency in mammalian cells from the original average 4.29% to 48.94% with an average 26.52-fold improvement, substantially outperforming the popular PAM-free version of nuclease-SpRY. Moreover, a single AAV delivery of eTasR *in vivo* effectively alleviates the disease phenotype of hereditary tyrosinemia type I (HT1) mouse model through targeted *Hpd* editing. Together, these findings establish eTasR as a compact, PAMless, and highly efficient genome editing tool with broad therapeutic and research applications.

## Introduction

The TIGR-Tas system, discovered in 2025 by Faure et al., defines a distinct family of RNA-guided DNA-targeting systems found across bacteriophages, archaeal viruses and parasitic bacteria.^1^ Two features make TIGR-Tas exceptionally attractive for genome engineering. First, TasR cleaves target DNA in a completely PAM-independent manner, distinguishing it from CRISPR-Cas nucleases and enabling access to genomic sites inaccessible to PAM-restricted editors. Second, the TasR effectors are very compact, with only ∼331 amino acids for TaTasR or 338 amino acids for ParTasR, making them highly amenable to delivery by adeno-associated virus (AAV) (Fig. 1a). Despite these advantages, wild-type ParTasR was reported to exhibit only 0.8–3.6% indel formation in mammalian cells, severely limiting its utility for basic research and therapeutic genome editing.^1^

**Fig. 1.**
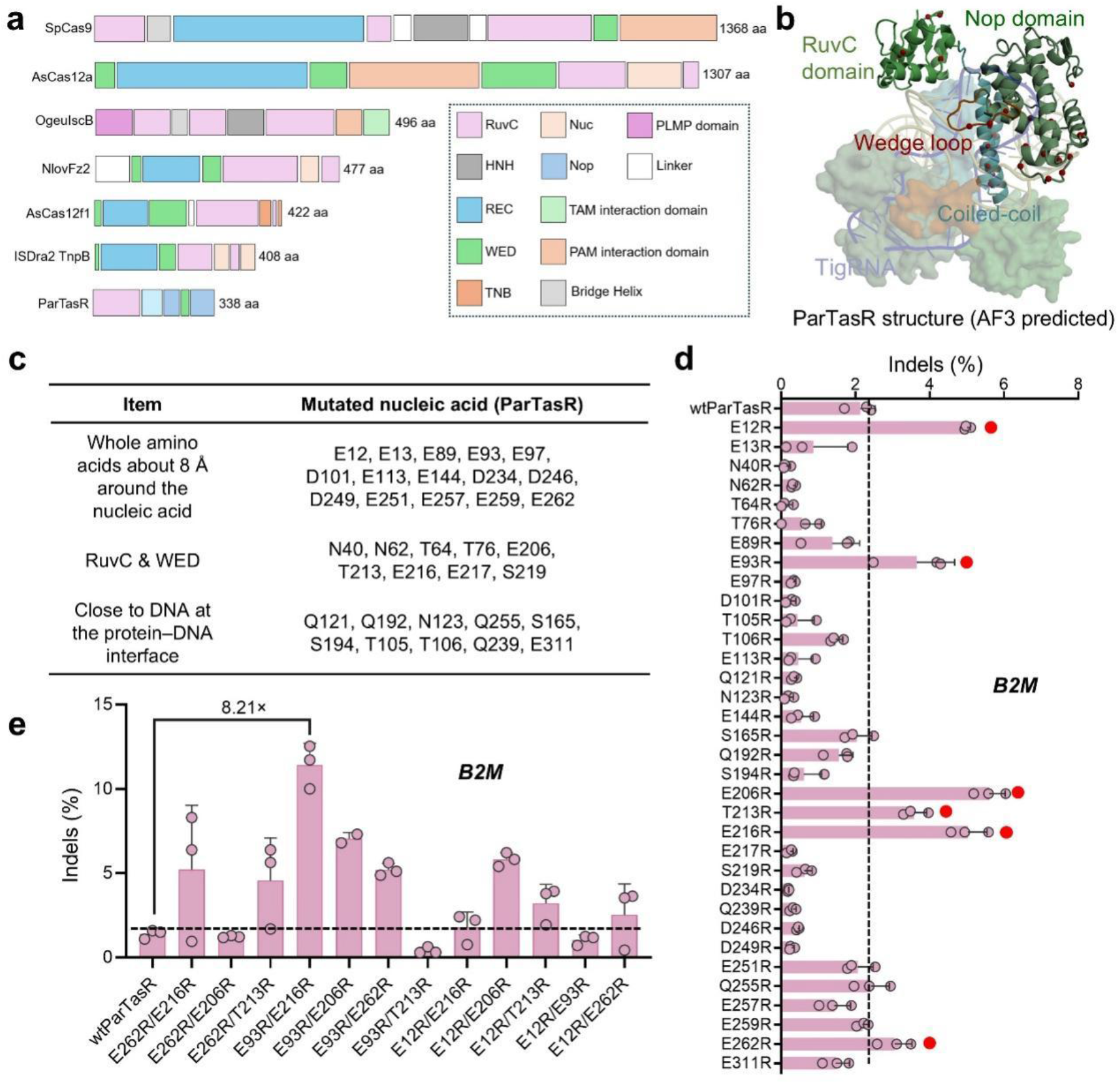
Structure-guided engineering of ParTasR for increased genome-editing efficiency. **a**, Comparison of the size of ParTasR with representative genome-editing endonucleases, including SpCas9, AsCas12a, OgeuIscB, NlovFz2, AsCas12a1, and ISDra2 TnpB. **b**, Predicted structure of the ParTasR-tigRNA-target DNA complex generated by AlphaFold 3. The following 34 candidate single-mutation amino acid residues are highlighted in red in the structure figure. **c**, Candidate ParTasR amino acid residues selected for arginine substitution based on structural analysis. Candidates were selected according to three criteria: (i) all non-positively charged residues within 8 Å of the nucleic acids; (ii) additional non-positively charged residues located within the RuvC or WED domains; and (iii) non-positively charged residues at the protein-target DNA interface. Candidate residues were individually substituted with arginine (R). **d**, Indel frequencies at the endogenous *B2M* locus generated by wild-type ParTasR and ParTasR variants bearing single amino acid substitutions. **e**, Indel frequencies at the endogenous *B2M* locus generated by ParTasR variants bearing two amino acid substitutions. For **d–e**, data are presented as mean ± s.d. from three independent experiments (*n* = 3).

The development of miniature RNA-guided nucleases has attracted considerable interest as a strategy to overcome the *in vivo* delivery constraints imposed by the large size of many Cas effectors. *Staphylococcus aureus* Cas9 (SaCas9, ∼1,053 aa) has been extensively explored for *in vivo* genome editing, but its stringent PAM requirement (NNGRRT) constrains its targeting scope.^2^ Type V Cas12f systems (∼400–700 aa) have been engineered to achieve robust mammalian editing through optimization of the protein and guide RNA.^3,4^ Engineered TnpB variants (∼400 aa), the evolutionary ancestors of Cas12, have achieved 23–42% indel frequencies across multiple loci through mutational scanning optimization.^5^ The OMEGA systems have likewise proven more amenable to engineering: structure-guided optimization of IscB (∼400–500 aa) has yielded variants with editing efficiencies exceeding 90% in human cells,^6^ and engineered Fanzor nucleases (∼400–700 aa) have achieved expanded TAM recognition and average indel frequencies above 70%.^7^ Despite these advances, compact RNA-guided nucleases generally remain constrained by PAM or TAM requirements that limit their targeting scope. Engineered PAMless Cas9 variants such as SpRY overcome this constraint but exhibit reduced editing activity in certain sequence contexts.^8,9^ Thus, a compact and highly efficient genome editor with PAM-independent targeting capability remains an unmet need. The recently discovered TIGR-Tas system,^1^ with its intrinsic PAM independence and ultra-compact architecture, therefore represents a promising starting point for developing such an editor.

In this study, we engineered ParTasR to develop a compact and highly efficient genome editor. Guided by the predicted structure of the TasR–tigRNA–DNA ternary complex, we identified a ParTasR variant carrying E93R and E216R that exhibited an average 2.55-fold improvement in editing efficiency over wild-type ParTasR. Co-expression of the human 3′→5′ exonuclease TREX2 with ParTasR(E93R/E216R) yielded ParTasR(E93R/E216R)-P2A-TREX2, termed eTasR, which exhibited an average 26.52-fold improvement in editing efficiency over wild-type ParTasR. Moreover, the efficiency of eTasR is substantially higher than that of SpRY, the only currently available PAM-free nuclease.^8,9^ Notably, single-AAV delivery enabled efficient *Hpd* disruption *in vivo*, resulting in robust phenotypic rescue in HT1 mice. Together, these findings establish eTasR as a compact, PAMless and highly efficient genome editor with broad potential for genome engineering and therapeutic applications.

## Results

### Engineering an enhanced-efficiency ParTasR in mammalian cells

Two TasR effectors, TaTasR and ParTasR, have been shown to mediate genome editing in mammalian cells, with ParTasR exhibiting higher activity but still reaching only 3.6% indel formation.^1^ We therefore selected ParTasR as the starting point for engineering. Since the structure of ParTasR has not been experimentally resolved, we used AlphaFold 3 to predict the structure of ParTasR in complex with tigRNA and target DNA to guide rational engineering. Given the moderate sequence identity (36.10%) between ParTasR and TaTasR, we used the experimentally determined cryo-EM structure of TaTasR as a structural reference for model selection (Fig. 1b and Extended Data Fig. 1). We designed mutations according to three structure-guided criteria: (1) all non-positively charged residues within 8 Å of the nucleic acid molecules, including tigRNA and target DNA; (2) additional non-positively charged residues located within the RuvC and WED domains, as mutations in these regions have been reported to enhance editing efficiency;^10–13^ and (3) non-positively charged residues directly adjacent to the target DNA at the protein–DNA interface. These amino acid residues were individually substituted with strongly positively charged ones (arginine, R), with the rationale that introducing positively charged side chains might favor interactions with the negatively charged nucleic acids and thereby enhance ParTasR activity.^14,15^

Using this strategy, we generated 34 ParTasR mutants (Fig. 1c). The mutants were co-transfected with a tigRNA plasmid targeting the endogenous human *B2M* locus (co-expressing EGFP) into HEK293 cells. At 72 h post-transfection, EGFP-positive cells were enriched by fluorescence-activated cell sorting (FACS; Supplementary Note), followed by genomic DNA extraction, PCR amplification of the target locus and high-throughput sequencing (HTS). The HTS results showed that six single mutants exhibited higher editing activity than wild-type ParTasR, including E12R and E93R located in the RuvC domain; E206R, T213R and E216R in the WED domain; and E262R in the Nop domain (Fig. 1d). Given that these efficiency-enhancing mutations were distributed across distinct functional domains, we speculated that combining mutations from different regions may produce synergistic improvement. We therefore constructed and evaluated a series of cross-domain double-mutant combinations. Among these variants, ParTasR(E93R/E216R), combining mutations in the RuvC and WED domains, showed the strongest synergistic enhancement, reaching 11.41% editing efficiency compared with 1.39% for wild-type ParTasR, corresponding to an 8.21-fold increase (Fig. 1e).

### Overexpression of the exonuclease TREX2 further enhances ParTasR editing efficiency

Since ParTasR cleavage generates DNA ends with 3′ overhangs, which can be directly re-ligated by cellular DNA repair pathways when the ends are compatible, the efficiency of indel formation is thereby limited.^16,17^ Based on this principle, we hypothesized that TREX2, by processing 3′ overhangs to prevent precise rejoining via cNHEJ (classical non-homologous end joining) toward mutagenic end joining, would effectively improve editing efficiency (Fig. 2a). Therefore, we selected the 3′→5′ exonuclease TREX2 (236 aa) and co-expressed it with wild-type ParTasR in HEK293T cells targeting five endogenous loci to evaluate its performance.^18–19^ The 5′→3′ exonuclease T5 exonuclease (T5 exo, 291 aa) was included as a control.^20^ The results showed that co-transfection of TREX2 consistently increased indel frequency (from the original 0.06-4.16% to 5.07-53.68%), whereas expression of T5 exo did not lead to obviously improved efficiency, which was consistent with expectations (Fig. 2b).

**Fig. 2.**
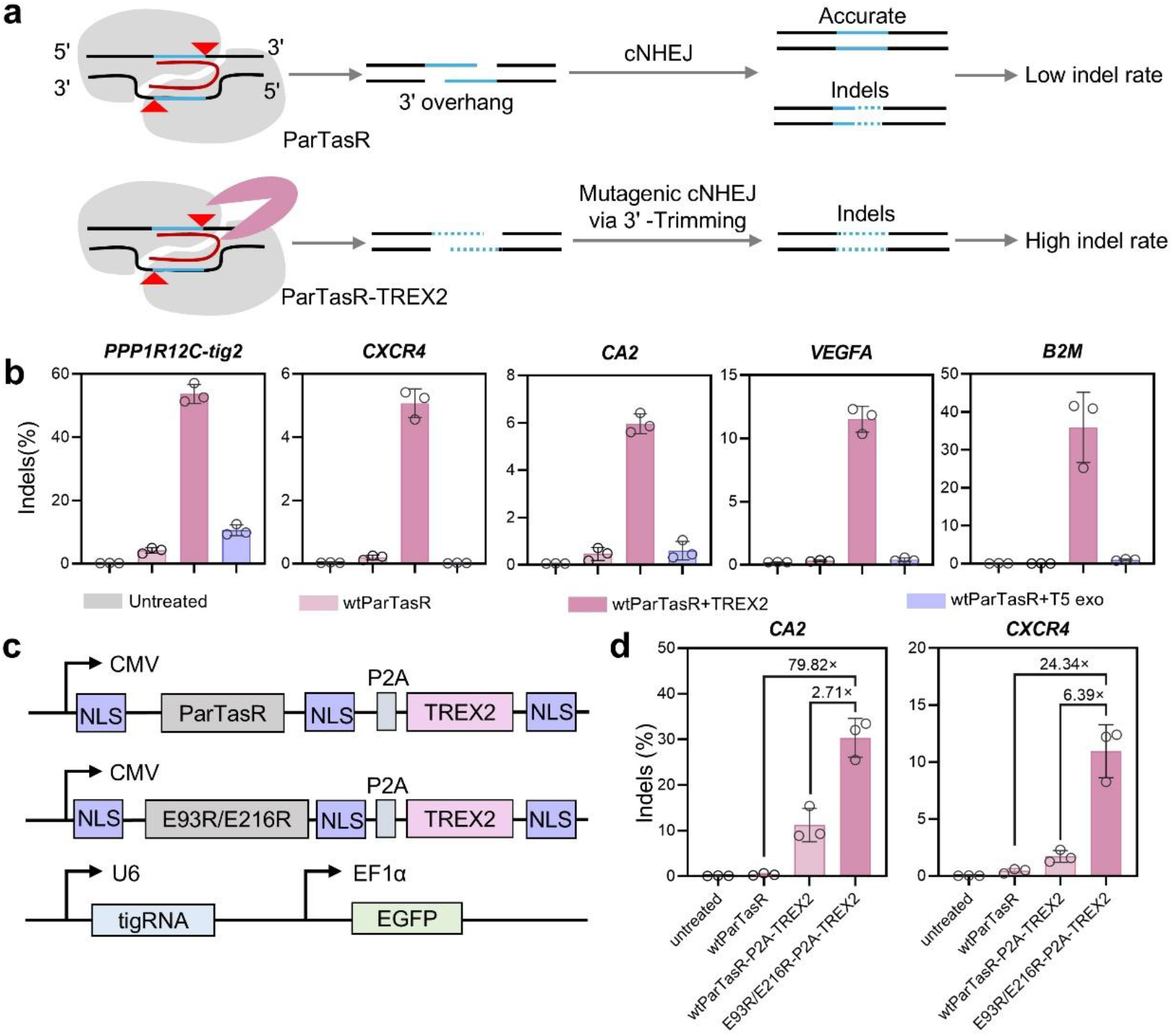
TREX2 enhances ParTasR-mediated genome editing in mammalian cells. **a**, Schematic illustrating the proposed mechanism by which TREX2 enhances ParTasR-mediated genome editing by processing 3′ DNA overhangs. cNHEJ, classical nonhomologous end joining. **b**, Effects of TREX2 or T5 exonuclease (T5 exo) on the editing efficiency of wild-type ParTasR at five endogenous genomic loci. **c**, Schematic of the P2A-linked bicistronic cassette for co-expression of wild-type ParTasR or ParTasR(E93R/E216R) with TREX2. **d**, Effects of co-expression of wild-type ParTasR or ParTasR(E93R/E216R) with TREX2 from a P2A-linked bicistronic cassette at two endogenous genomic loci. For **b** and **d**, the data are presented as mean ± s.d. from three independent experiments (*n* = 3).

We next applied this strategy to ParTasR(E93R/E216R) to test whether TREX2 could further enhance its editing efficiency at two endogenous loci. The results showed that co-expression of TREX2 with ParTasR(E93R/E216R) increased editing efficiency relative to wild-type ParTasR co-expressed with TREX2, with the largest increase observed at *CXCR4* (up to 6.39-fold). Relative to wild-type ParTasR, co-expression of TREX2 with ParTasR(E93R/E216R) increased editing efficiency by up to 79.82-fold, with the largest increase observed at *CA2* (Fig. 2c and Fig. 2d). We therefore selected ParTasR(E93R/E216R)-P2A-TREX2, hereafter termed enhanced ParTasR (eTasR), for further investigation.

### eTasR exhibits efficient and specific genome editing

To further unbiasedly profile the characteristic of eTasR, we selected 13 additional endogenous targets to evaluate its performance against SpRY.^8^ HEK293T cells were co-transfected with the indicated ParTasR variants or SpRY together with their corresponding tigRNA or sgRNA, respectively. After 72 h, EGFP-positive cells were sorted and target editing efficiency was evaluated by HTS. Analysis of HTS results revealed that the editing efficiencies of eTasR and ParTasR(E93R/E216R) were 7.26-91.32% and 0.29%–34.13%, respectively, representing an average increase of 26.52 fold and 2.55-fold compared to wild-type ParTasR with efficiencies 0.30-12.43%. More importantly, eTasR demonstrated substantially higher editing efficiency than SpRY across 12 of the 13 tested target sites. ParTasR(E93R/E216R) showed higher editing efficiencies than SpRY at five target sites, lower efficiencies at six target sites, and comparable efficiencies at the remaining two target sites (Fig. 3a). Overall, eTasR exhibited significantly higher editing efficiency than ParTasR (E93R/E216R), wild-type ParTasR, and SpRY. Meanwhile, ParTasR (E93R/E216R) demonstrated significantly improved efficiency relative to wild-type ParTasR, yet remained comparable to SpRY (Fig. 3b). We also measured the editing efficiency of eTasR without FACS enrichment. Notably, eTasR achieved 25.93–74.24% indel frequencies across multiple target sites without FACS (Extended Data Fig. 2).

**Fig. 3.**
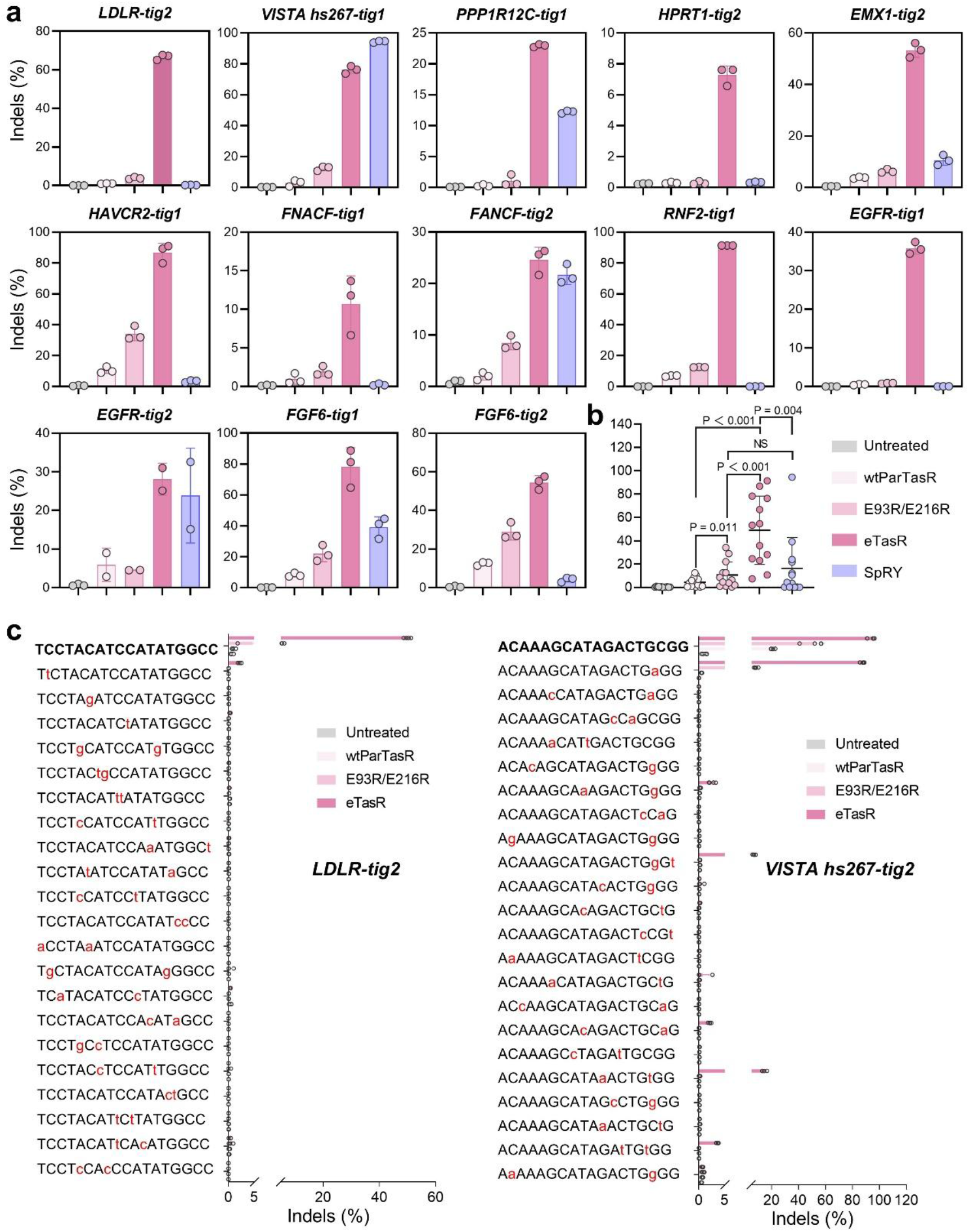
eTasR exhibits efficient and specific genome editing. **a**, Comparison of indel frequencies generated by wild-type ParTasR, ParTasR(E93R/E216R), eTasR and SpRY across 13 endogenous genomic loci in HEK293T cells. Data are presented as mean ± s.d. from three independent experiments (*n* = 3). **b**, Summary of the editing activities of wild-type ParTasR, ParTasR(E93R/E216R), eTasR and SpRY across the tested endogenous loci. Each dot represents the mean indel frequency from three independent biological replicates at one endogenous locus. *P* values were determined using a two-tailed paired *t*-test. NS, not significant. Data are presented as mean ± s.d. **c**, Analysis of off-target editing by ParTasR(E93R/E216R) and eTasR at *LDLR-tig2* and *VISTA hs267-tig2*. The top sequence in each panel represents the on-target sequence, and the sequences below represent Cas-OFFinder-predicted off-target sites. Mismatched bases relative to the on-target sequence are highlighted in red. Data are presented as mean ± s.d. from four independent experiments (*n* = 4).

To further evaluate the specificity of eTasR and ParTasR(E93R/E216R), we used Cas-OFFinder to predict potential off-target sites for *LDLR-tig2* and *VISTA hs267-tig2* based on sequence similarity to the corresponding tigRNAs^21^. HTS analysis of 43 predicted off-target sites revealed no detectable off-target editing at the vast majority of sites for either eTasR or ParTasR(E93R/E216R). Seven off-target sites showed detectable editing, six of which exhibited low editing frequencies, and only one tigRNA containing a single mismatch exhibited high off-target editing (Fig. 3c). It is speculated that the cause of off-target is related to the selection of tigRNA. Further optimization of tigRNA design may improve the specificity of eTasR. Together, these results suggest that eTasR is an efficient and specific genome editing tool.

### Single-AAV delivery of eTasR enables efficient gene therapy for HT1 *in vivo*

Owing to its compact size, eTasR can be co-packaged with its cognate tigRNA into a single recombinant adeno-associated virus (rAAV) vector, positioning it as a promising therapeutic candidate for monogenic disorders,^22,23^ including hereditary tyrosinemia type I (HT1). We next tested whether eTasR could rescue the disease phenotype in a mouse model of HT1 by editing *Hpd* gene, which is involved in the metabolic pathway affected in HT1.^24^ We therefore packaged eTasR together with a tigRNA targeting *Hpd* exon 3 into a single AAV8 vector and administered it intravenously to *Fah^−/−^* HT1 mice via tail-vein (Fig. 4a). Two weeks after AAV8 injection, nitisinone (NTBC) was withdrawn, and body weight was monitored for 30 days following NTBC withdrawal (Fig. 4b). Whereas PBS-treated control mice exhibited progressive body weight loss exceeding 20%, AAV-treated mice maintained stable weight gain following NTBC withdrawal (Fig. 4c). On day 30 after NTBC withdrawal, blood was collected from AAV-treated mice for serum isolation. The mice were then euthanized for liver genomic DNA extraction and assessment of *Hpd* editing efficiency, as well as for collection of liver tissues for histological and pathological analyses.

**Fig. 4.**
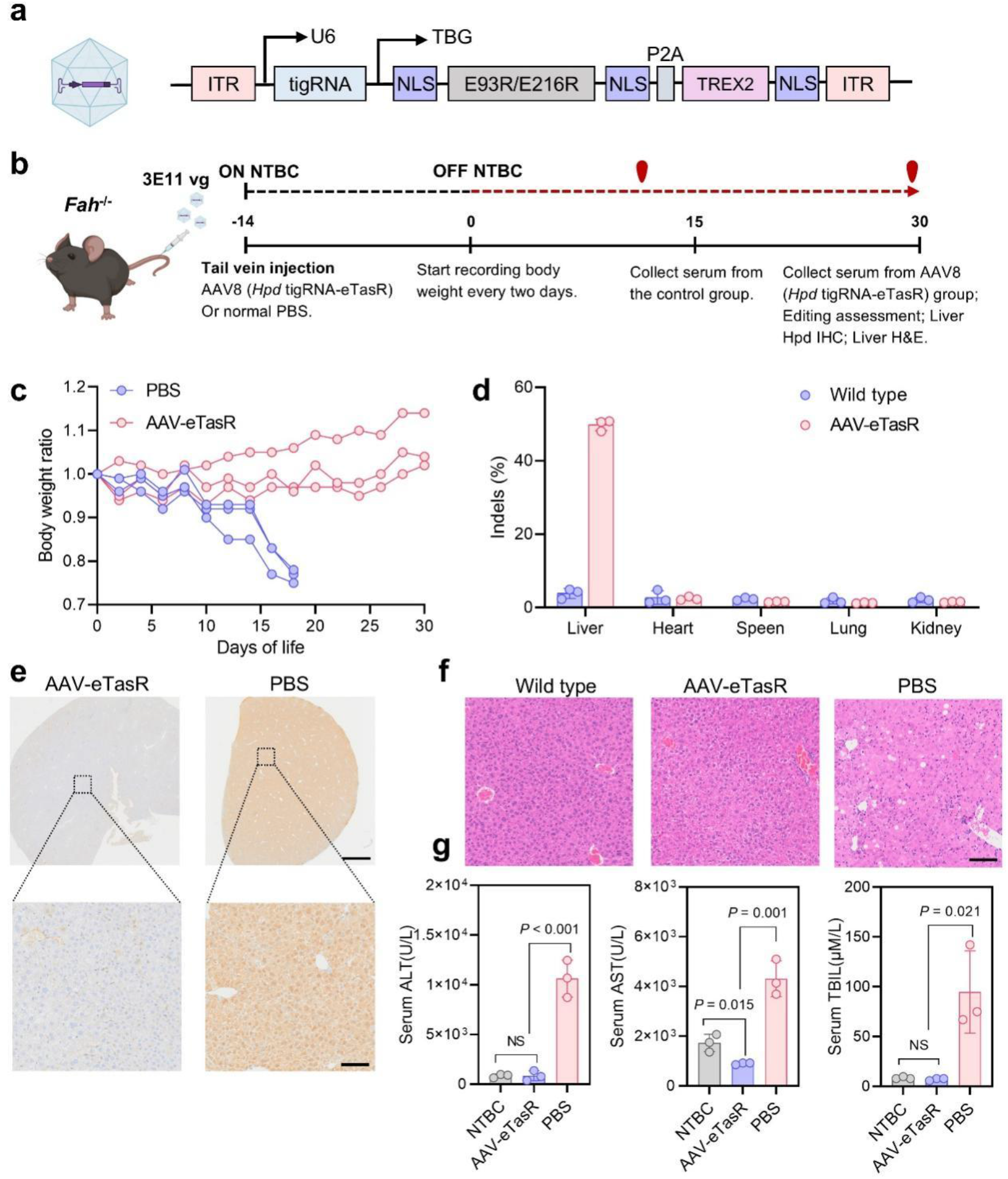
eTasR enables efficient gene therapy for HT1 following single-AAV delivery. **a**, Schematic of the single-AAV8 vector encoding eTasR and the *Hpd*-targeting tigRNA. **b**, Nitisinone (NTBC) was withdrawn 2 weeks after AAV administration, and mice were monitored for 30 days following NTBC withdrawal. Organs and serum were collected from AAV-treated mice at day 30 and from PBS-treated control mice at day 18 after NTBC withdrawal. **c**, Body weight trajectories following NTBC withdrawal. Body weight on the day of NTBC withdrawal (day 0) was normalized to the baseline for each mouse. **d**, *Hpd* indel frequencies in the liver, heart, kidney, lungs and spleen of PBS and AAV injected mice. **e**, Immunohistochemical (IHC) staining of Hpd protein in liver sections from AAV-treated mice at day 30 and PBS-treated control mice at day 18 after NTBC withdrawal. Scale bars, 1 mm (top) and 100 μm (bottom). **f**, Hematoxylin and eosin (H&E) staining of liver tissue sections from wild-type mice, AAV injected mice and PBS treated control mice at day 18 after NTBC withdrawal. Scale bars, 100 μm. **g**, Serum ALT, AST, and TBIL levels in peripheral blood from NTBC-treated mice, AAV injected mice at day 30 and PBS-treated mice at day 18. ALT, alanine aminotransferase; AST, aspartate aminotransferase; TBIL, total bilirubin. Statistical significance was determined using unpaired two-tailed Student’s *t*-tests. NS, not significant. For **d** and **g**, data are presented as mean ± s.d. from three independent mouse (*n* = 3).

The HTS results showed that *Hpd* indel frequency reached 49.87% in the liver of AAV-treated mice, whereas no appreciable editing was detected in the heart, kidney, lung or spleen (Fig. 4d). Consistent with the genomic editing results, immunohistochemical (IHC) analysis showed reduced Hpd protein expression in the liver of AAV-treated mice (Fig. 4e). Hematoxylin and eosin (H&E) staining further showed that liver architecture was largely preserved in AAV-treated mice, with no apparent inflammatory infiltration, necrosis or fibrosis (Fig. 4f). Serum biochemical analyses demonstrated that alanine aminotransferase (ALT), aspartate aminotransferase (AST) and total bilirubin (TBIL) levels in AAV-treated mice were comparable to those in wild-type mice, whereas PBS-treated HT1 mice exhibited marked elevations in serum transaminases (Fig. 4g). These findings indicate that eTasR can effectively rescue the disease phenotype of HTI when delivered *in vivo* via a single AAV vector, suggesting that eTasR holds promising potential for treating genetic disorders in the future.

## Discussion

In recent years, development of efficient small nucleases has become a focal point in the field due to their compact size and ease of *in vivo* delivery.^25–26^ In this study, we engineered ParTasR to develop an effective, PAM-free, and compact genome editor system. Through structure-guided rational design, we obtained ParTasR(E93R/E216R), and further enhanced its efficiency by co-expressing TREX2 to generate eTasR, achieving average improvements of 2.55-fold and 26.52-fold compared to wild-type ParTasR, respectively—representing a significant technological advancement. Notably, across the tested loci, eTasR achieved editing efficiencies of up to 91.32%, compared with the previously reported maximum of 3.6% for wild-type ParTasR, surpassing or matching the performance of existing small nucleases, such as SaCas9, Cas12f, TnpB, IscB, and Fanzor^2–7^. It thus effectively transforms previously inefficient or poorly functional tools into ones suitable for broad applications. More importantly, ParTasR’s intrinsic PAM independence confers a distinct practical advantage over other small nucleases.

In addition to achieving performance comparable to small nucleases, eTasR has also demonstrated superior results in head-to-head comparisons with SpRY, the recently reported optimal PAM-independent SpCas9 mutant.^8^ However, since the editing efficiency of PAM-independent SpRY remains lower than that of the classic SpCas9 with NGG PAM, which is widely recognized as a benchmark in both basic research and clinical disease therapy, further improvement of eTasR to surpass or match SpCas9 remains a key future goal. Moreover, while eTasR shows enhanced editing efficiency compared to wild-type ParTasR, off-target editing was detectable at a subset of predicted sites, indicating that future efforts must focus on improving overall performance while minimizing off-target activity.

Derivative genome editors based on CRISPR/Cas9, such as base editors and prime editors, have also emerged as next-generation tools in the field of gene editing due to their unique advantages. ^27–31^ Given eTasR’s PAM independence and compact size, future development of eTasR-based derivative editing tools incorporating deaminases or reverse transcriptase may surpass a range of existing CRISPR/Cas9-based tools, representing an important direction for advancement.

In summary, we engineered ParTasR to develop an efficient, PAM-free, and compact genome editing system—eTasR—and demonstrated its value in genome editing and *in vivo* therapeutic applications. In the future, eTasR holds great potential for broader applications in various research areas, including disease therapy, large animal model development, genetic screening, and agricultural applications.

## Methods

### Structural prediction and residue selection for ParTasR engineering

The three-dimensional structure of the ParTasR-tigRNA-DNA complex was predicted using the AlphaFold 3 Server (https://alphafoldserver.com) following the previously described protocol.^32^ The prediction inputs included two copies of ParTasR (338 amino acids each), one tigRNA (36 nt; 5′–AGCCAUGCAAGCCCUGAAACCCAUUGCCUUAGUGCG–3′), and one target DNA duplex (38 bp; top strand 5′–GTGGTTTCGATGCAAGCCCCTAAGGCAATCATCAATTT–3′, bottom strand 5′–AAATTGATGATTGCCTTAGGGGCTTGCATCGAAACCAC–3′).^1^ Four Mg²⁺ ions were included and all other parameters were kept at the server default settings.

Among the returned models, the one that most closely recapitulated the overall architecture of the experimentally determined TasR complex (PDB: 9MTY) was selected for downstream analysis. Candidate residues were identified using PyMOL by selecting ParTasR residues with any heavy atom within 8 Å of a nucleic acid heavy atom (either target DNA or tigRNA). Residues that were already arginine were excluded. All remaining candidates were substituted with arginine and subsequently evaluated for editing efficiency. The amino acid sequences of all mutants are provided in Supplementary Table 1.

### Sequence alignment

The amino acid sequences of TaTasR and ParTasR were aligned using the ClustalW online server (https://www.genome.jp/tools-bin/clustalw). The resulting multiple sequence alignment was visualized using the ESPript 3.2 web server (https://espript.ibcp.fr/ESPript/) to highlight conserved residues and secondary structure elements. Furthermore, the pairwise percentage identity between the two sequences was calculated and analyzed using Jalview 2.11.5.2 (https://www.jalview.org/).

### Plasmid construction

The human codon-optimized coding sequence of ParTasR was synthesized by Genewiz (Suzhou, China). To generate ParTasR mutants, DNA fragments were amplified by PCR using KOD-Plus-Neo DNA polymerase (TOYOBO, Code: KOD-401) and assembled into linearized vectors using the ClonExpress MultiS One Step Cloning Kit (Vazyme) following the manufacturer’s instructions. All constructed plasmids were verified by Sanger sequencing.

For tigRNA and sgRNA expression vectors, complementary oligonucleotides (Supplementary Table 2) corresponding to the target sequences were annealed by heating at 95 °C for 5 min and gradually cooling to room temperature. The annealed duplexes were ligated into BbsI-linearized U6-tigRNA-EF1α-EGFP or U6-sgRNA-EF1α-EGFP vectors using T4 DNA ligase (Thermo Fisher Scientific). All insertions were confirmed by Sanger sequencing.

### Cell culture, transfection, and fluorescence-activated cell sorting

HEK293T cells (ATCC, CRL-3216) were cultured in Dulbecco’s Modified Eagle Medium (DMEM, Thermo Fisher Scientific) supplemented with 10% fetal bovine serum (FBS, Gibco) and 1% penicillin-streptomycin (Thermo Fisher Scientific). All cells were maintained at 37 °C in a humidified atmosphere containing 5% CO_2_. For gene editing experiments, HEK293T cells were seeded into 24-well plates and cultured until reaching approximately 70–80% confluence prior to transient transfection. Each well was transfected with 1 μg of plasmid DNA using polyethylenimine (PEI, Polysciences). The transfection mixture contained 750 ng of editor expression plasmid, 250 ng of tigRNA or sgRNA expression plasmid, and 3 µL of PEI, which were diluted separately in 50 µL of DMEM, mixed thoroughly, and incubated at room temperature for 15 min before addition to the cells.

To assess the editing activity, cells were harvested 72 h post-transfection. Briefly, cells were washed with phosphate-buffered saline (PBS), trypsinized with 0.25% trypsin-EDTA, resuspended in DMEM, and filtered through 40 μm cell strainers to obtain single-cell suspensions. Fluorescent positive cells were isolated using fluorescence-activated cell sorting (FACS). Genomic DNA was extracted from 20,000 sorted cells using the QuickExtract™ DNA Extraction Solution (Lucigen) according to the manufacturer’s instructions.

### High-throughput sequencing and data analysis

Target genomic regions and potential off-target loci were amplified by PCR for deep sequencing analysis, the corresponding target sites, off-target sites, and amplification primers are listed in Supplementary Table 2. To prepare the high-throughput sequencing libraries, primary PCR was performed using 2× Ultra-Rapid Hot Start PCR Master Mix (with Dye) with primers containing 5′ adapter sequences (forward: 5′-GGAGTGAGTACGGTGTGC-3′; reverse: 5′-GAGTTGGATGCTGGATGG-3′). Subsequently, a second round of PCR was performed using 2× Taq Master Mix with barcoded primers. The barcoded PCR products were pooled and subjected to sequencing on an Illumina HiSeq platform (Annoroad Gene Technology). FASTQ files were processed to obtain full-length sequences as previously described.^33^ Editing efficiencies were calculated using CRISPResso2 software.^33^

### In silico prediction of tigRNA off-target sites

Potential genomic off-target sites for each tigRNA spacer were predicted genome-wide using Cas-OFFinder v3.0.0b3.^21^ Searches were performed against the human reference genome hg38, allowing up to three mismatches and no DNA or RNA bulges, without imposing a PAM constraint. Mismatch positions along each aligned DNA site were parsed from Cas-OFFinder output.^21^

### AAV package and treatment

Mouse feeding conditions were as described above. Recombinant adeno-associated virus (AAV) carrying the U6-Hpd tigRNA-TBG-eTasR cassette was packaged and titrated by Packgene Co., Ltd. (Guangzhou, China). A total of 3E11 vector genomes (vg) of AAV was administered to 6–8-week-old *Fah*^-/-^ mice via tail vein injection. The control group received an equivalent volume of PBS.

### Immunohistochemistry and histology

Liver tissues were collected and processed for immunohistochemistry (IHC) and hematoxylin and eosin (H&E) staining by Wuhan Service Biotechnology Co., Ltd. (Wuhan, China). For IHC and H&E staining, mouse livers were fixed with 4% paraformaldehyde (PFA), embedded in paraffin, and sectioned at 4 μm thickness. Anti-HPD monoclonal antibody (67552-1-Ig, 1:1000) was used for IHC to detect Hpd-positive hepatocytes.

### Serum analysis

To minimize batch effects, serum samples collected at different time points within the same experiment were analyzed concurrently. Levels of alanine aminotransferase (ALT), aspartate aminotransferase (AST), and total bilirubin (TBIL) were measured using assay kits from Solarbio (Beijing, China; cat. no. BC1555, BC1565, and BC6454, respectively). All procedures were performed according to the manufacturer’s instructions.

### Statistics and reproducibility

Data are presented as mean ± SD from three independent biological replicates performed on different days. All graphs were generated using GraphPad Prism 8.0.2 (GraphPad Software, San Diego, CA, USA). Statistical significance between two groups was assessed using two-tailed Student’s t-test (GraphPad Prism 8.0.2). The specific statistical tests used are indicated in the corresponding figure legends. P value < 0.05 was considered statistically significant.

## Data availability

All data supporting the findings of this study are available within the article and its Supplementary Information. The raw high-throughput sequencing data generated in this study have been deposited to the NCBI Sequence Read Archive (SRA) under accession code PRJNA1524974. Plasmids encoding ParTasR(E93R/E216R) and eTasR will be available from WeKwikGene plasmid repository. Source data are provided with this paper.

## Author contributions

X.Z. conceived and supervised the project. Q.Z. designed the project. Q.Z., Z.Z., and Q.C. performed the experiments and data analysis with assistance from X.M., Y.Z., and M. S.. X.Z. and Q.Z. wrote the manuscript with input from all the authors.

**Correspondence and requests for materials** should be addressed to X.Z.

