## Supplemental information for "Engineered Miniature ParTasR Enables Efficient PAMless Genome Editing"

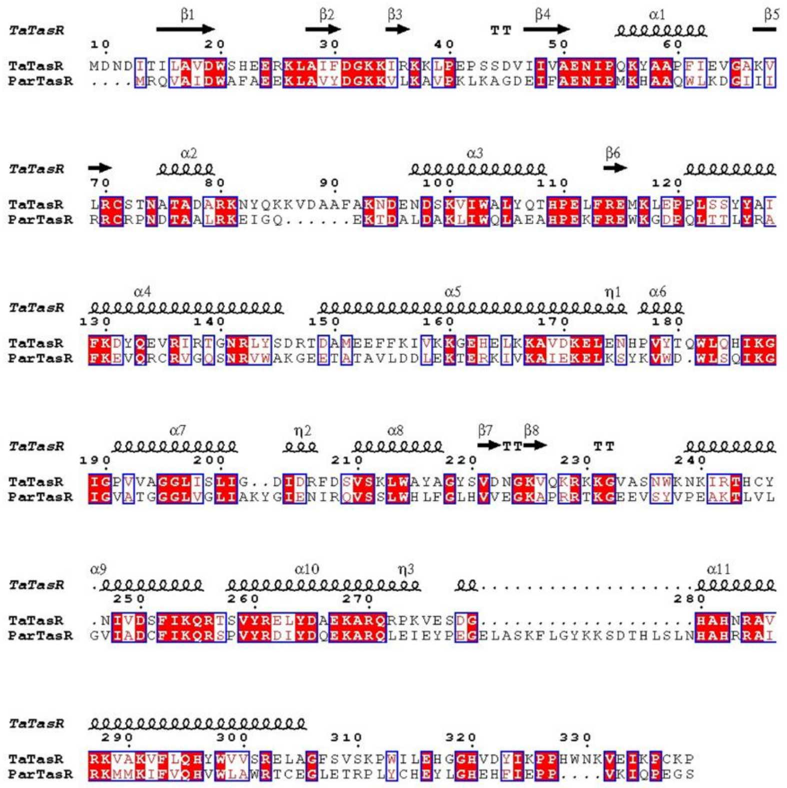

**Extended Data Fig.1 |** Sequence alignment of ParTasR and its ortholog TaTasR（9MTY）

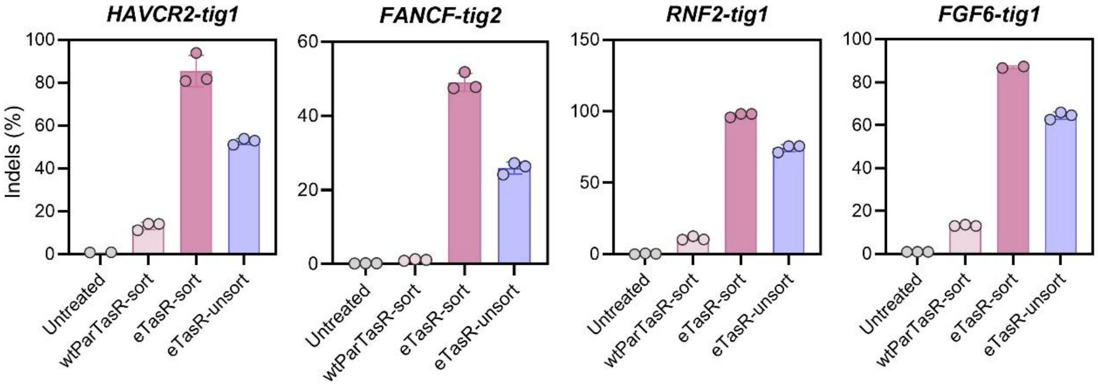


**Extended Data Fig.2 |** Comparison of eTasR-mediated editing efficiency under sorted and unsorted conditions at four endogenous genome loci. The data are presented as the mean ± s.d. from three independent experiments (n = 3).

### Supplementary note

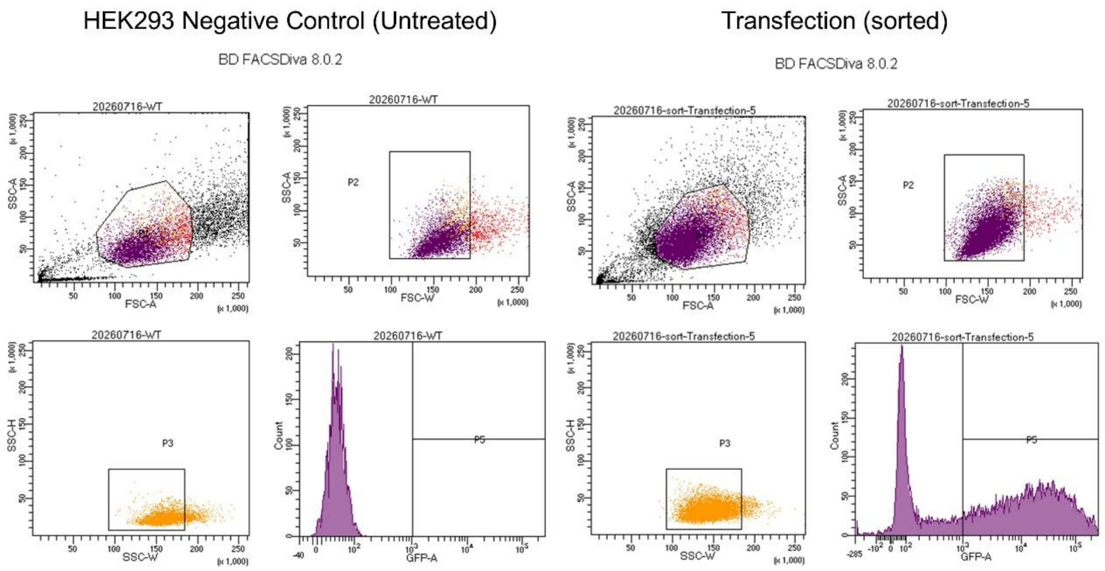


Tube: WT

| Population | #Events | %Parent | %Total |
|---|---|---|---|
| All Events | 10,000 | #### | 100.0 |
| P1 | 4,567 | 45.7 | 45.7 |
| P2 | 3,935 | 86.2 | 39.4 |
| P3 | 3,429 | 87.1 | 34.3 |
| P5 | 0 | 0.0 | 0.0 |

Tube: sort-Transfection-5

| Population | #Events | %Parent | %Total |
|---|---|---|---|
| All Events | 14,441 | #### | 100.0 |
| P1 | 10,721 | 74.2 | 74.2 |
| P2 | 10,410 | 97.1 | 72.1 |
| P3 | 10,043 | 96.5 | 69.5 |
| P5 | 6,144 | 61.2 | 42.5 |
